# Organ Scaffold Characterizations Using Intravital Microscopy

**DOI:** 10.1101/2022.01.20.477052

**Authors:** Peter R. Corridon, Anousha A. Khan

## Abstract

This study contains intravital microscopy (IVM) data to characterize organ scaffolds created using decellularization. Decellularization creates cell-free collagen-based matrices from native organs, which can be used as scaffolds for regenerative medicine applications. This data set contains *in vivo* assays evaluating the effectiveness of decellularization and structural and functional integrity of the acellular nephron in the post-transplantation environment. Qualitative and quantitative assessments of scaffold DNA concentrations, tissue fluorescence signals, structural and functional integrities of various decellularized nephron segments, and velocity within the microcirculation were acquired and compared to the native (non-transplanted) organ. Cohorts of 2-3-month-old male Sprague Dawley rats were used: non-transplanted (n = 4), transplanted day 0 (n=4), transplanted day 1 (n = 4), transplanted day 2 (n = 4), and transplanted day 7 (n = 4). OIF formatted files illustrate IVM measurement processes and are made publicly available in a data repository. Such data is intended to support scientific reproducibility, complement future renal studies, and extend the use of this powerful imaging application to analyze other organ scaffold systems.

|  |  |
| --- | --- |
| Measurements(s) | DNA quantification • tissue fluorescence • microvascular leakage • tubular and glomerular integrity • renal capillary blood velocity |
| Technology type(s) | intravital microscopy • multiphoton microscopy • UV-visible spectroscopy |
| Sample Characterization(s) | rats • native and decellularized kidneys |

Machine-accessible metadata file describing the reported data: https://doi.org/10.6084/m9.figshare.18703949.

## Background and Summary

The global rise in organ failure and scarcity of transplantable organs has heightened the quest for clinical alternatives. Within the past decade, advances in whole organ bioengineering technologies have demonstrated the potential to create such options. Decellularization is at the forefront of these technologies, as this method separates the extracellular matrix (ECM) from the native cells^1^. The ECM, with its preserved embedded structural and functional cues, can be employed as a natural scaffold to produce functioning replacement organs^2^. Such templates have been created from individual segments like blood vessels^3^ and tracheas^4^, and more complicated structures, including the lung, liver, and kidney^5^ using animal and human models.

Presently, the outcomes used to identify the efficacy of the decellularization process rely on the absence of cellular or nuclear remnants from the well-preserved tissue structure and maintenance of mechanical properties like native tissue. The protocol’s effectiveness also depends upon the size of the original organ, the concentration of the detergent, and the perfusion rate. For instance, several reports have shown how ionic detergents like sodium dodecyl sulfate (SDS) can routinely remove cells and genetic material from native structures quickly and effectively to produce homogeneous acellular renal templates^1,6–10^. These acellular scaffolds, primarily those made from solid organs, are often created from perfusion with this single surfactant or a blend of ionic and non-ionic surfactants. Nevertheless, a significant challenge to this process lies within the decellularized vasculature of complex organs like the kidney. The ideal conditions that will support scaffold longevity within transplantation environments are still unknown, as there is a limited understanding of decellularized vascular dynamics *in vivo*^11^.

To this end, a variety of macroscopic and microscopic approaches have been used to evaluate whole kidney scaffold structure and function^8–15^. In several animal models, such techniques have shown that the decellularization process successfully removes native cellular/tissue components and decellularizing agents while retaining the vascular and ECM structures. At the microscopic level, the vascular architecture must be preserved because it serves as an effective blueprint for cellular repopulation and an essential conduit to maintain organ viability post-transplantation. Furthermore, the efficient removal of scaffold remnants is critical for preventing negative biological and immunological consequences during recellularization and following transplantation. On the other hand, these approaches often limit the ability to perform investigations at the required spatial and temporal microscopic resolutions, highlighting the need for new ways to investigate these processes.

Interestingly, intravital microscopy (IVM) provides a unique opportunity to assess live morphological and functional aspects inside the kidney^17–20^. Consequently, the objective of this research was to obtain real-time *in vivo* characterizations of the decellularized kidney microarchitecture after transplantation using IVM. Techniques from previously refined methods that protect the existing vascular network while limiting cell toxicity were applied to generate acellular scaffolds^12, 21^. In this case, intact rat kidneys were extracted and decellularized with perfusion of SDS. The resulting scaffolds were orthotopically implanted into their respective donors. The removal rate of cellular/nuclear remnants and the viability of the decellularized network were then confirmed in real-time using nuclear and vascular dyes, respectively.

This data set also contains results from *in vivo* assays that evaluated the effectiveness of the decellularization process and structural and functional integrity of the acellular nephron in the post-transplantation environment. Qualitative and quantitative assessments of tissue autofluorescence, velocity within the microcirculation, and microarchitecture porosity of scaffolds were acquired in live rats on day 0, day 1, day 2, and day 7 after transplantation and compared to the native (non-transplanted) organs. This sophisticated imaging technology’s ability to evaluate renal dynamics *in vivo* is documented extensively^12–28^. Our manuscript aims to highlight methods of monitoring cellular/subcellular processes in real-time within transplanted kidney scaffolds, which can be extended to other organ scaffold systems. An overview of the experimental approach can be found in Fig. 1.

**Fig. 1.**
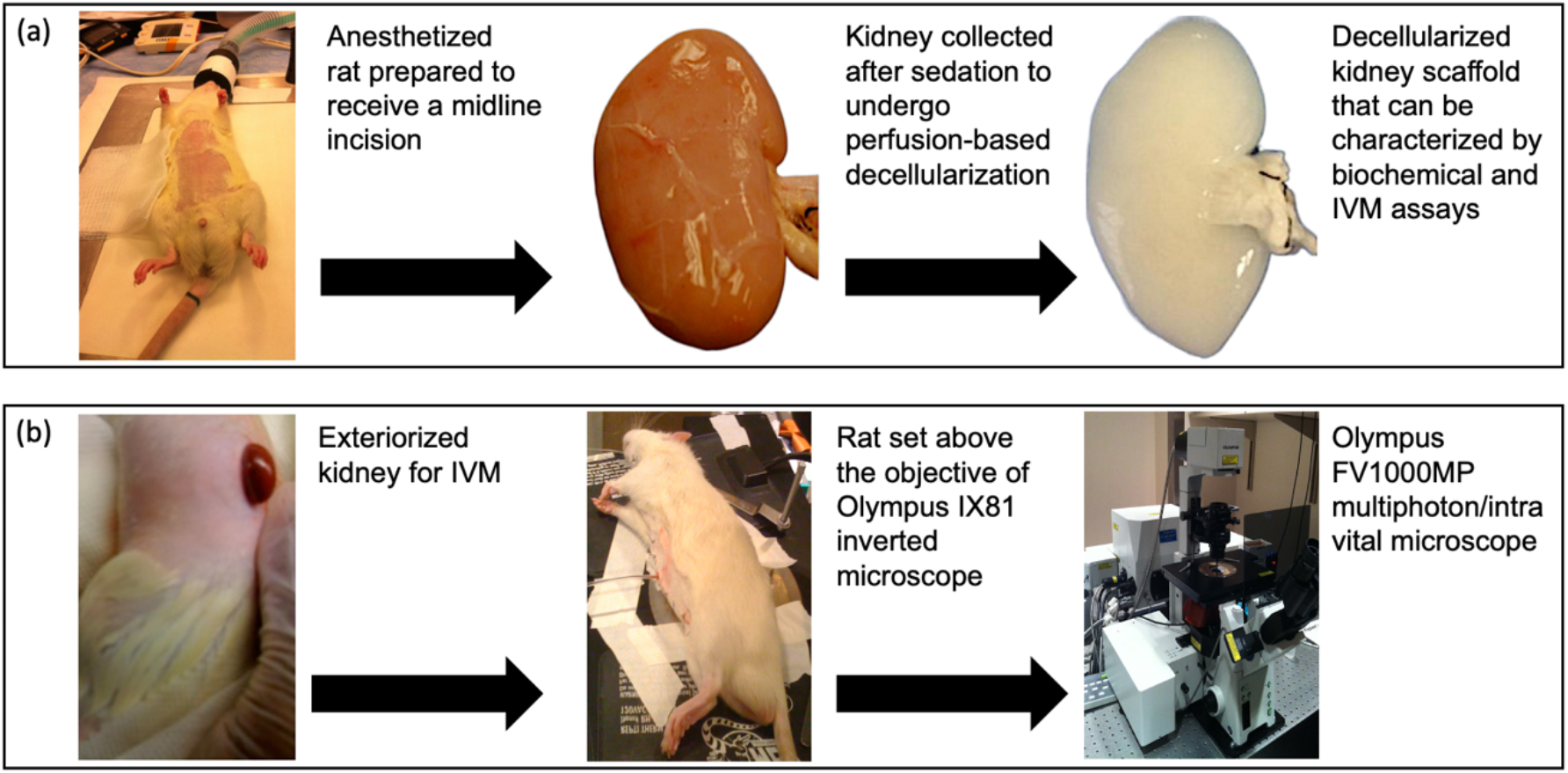
Overview of the study design. Photographs taken to illustrate (**a**) animal and organ manipulations to create acellular scaffolds for characterizations, (**b**) animal and organ preparations for IVM data acquisition and analysis.

## Methods

### Study design

This manuscript presents data obtained from a times series study used to characterize acellular kidney scaffolds using IVM. Experiments were conducted on twenty 2-3-month-old male Sprague Dawley rats. Each animal was randomly assigned to one of the following cohorts: non-transplanted (n = 4; n = 1 day 0, n = 1 day 1, n = 1 day 2, and n = 1 day 7), transplanted day 0 (n = 4), transplanted day 1 (n = 4), transplanted day 2 (n = 4), and transplanted day 7 (n = 4). An additional group of 4 rats was used for technical validation of the decellularization process, whereby both kidneys from each animal were used for biochemical assays to evaluate DNA and SDS concentrations in native (n = 4) and decellularized (n = 4) kidneys. The animals from that group received bilateral nephrectomies and were sacrificed after this procedure. The remaining sixteen animals that received unilateral nephrectomies were tracked by drawing lines around their tails with permanent markers and recording their associated cage numbers. Animal identifies were also duplicated in cage cards and laboratory records. The experimenters were not blinded to the animal’s identity throughout the entire experimental and analysis periods, and no animal was excluded from this study.

At the start of the study, all nephrectomies were performed to obtain whole kidneys that could be used to create the decellularized scaffolds. Transplantations and IVM studies were conducted after the kidneys were decellularized and the animals had enough time to recover from the radical left nephrectomy. The biochemical assays were performed during the recovery period to evaluate DNA and SDS contents. All experiments were performed in accordance with the Institutional Animal Care and Use Committee at Wake Forest University’s School of Medicine (Winston-Salem, NC, USA), the Animal Research Oversight Committee at Khalifa University of Science and Technology (Abu Dhabi, UAE), and ARRIVE criteria. These experimental procedures are outlined in detail below.

### Nephrectomies

Sham, unilateral, and bilateral nephrectomies were performed on 2-3-month-old male Sprague Dawley rats (Envigo, Indianapolis, IN) g. These animals were initially exposed to isoflurane (Webster Veterinary Supply, Devens, MA, USA) and then injected with 50 mg/kg pentobarbital intraperitoneally and placed on a heating pad to regulate core body temperature, shown in Fig. 1a. After that, midline incisions were then created along the aseptic regions, defined by the topical application of Betadine Surgical Scrub (Purdue Products L.P., Stamford, CT) to expose the left kidney. The corresponding renal artery, renal vein, and ureter were ligated with 4-0 silk (Fine Science Tools, Foster City, CA, USA). The native kidneys were extracted with intact renal arteries, veins, and ureters, as shown in Fig. 1a. The incision was closed in animals that received unilateral nephrectomies, and all these animals were given two weeks to recover. Similarly, midlines incisions and organ collection processes were performed during bilateral and sham nephrectomies.

### Scaffold decellularization, sterilization, and storage

The artery of each extracted kidney was connected to a 14-gage cannula (Fine Science Tools, Foster City, CA, USA) and PE-50 polyethylene catheter tubing (Clay Adams, Division of Becton Dickson, Parsippany, NJ) through which heparinized PBS (0.5-1 mL) was immediately perfused. The organs were then immersed in PBS, and the cannulated renal artery was connected to a peristaltic pump (Cole-Palmer, Vernon Hills, IL, USA). The kidneys were then perfused with 0.5% SDS (Sigma-Aldrich, St. Louis, MO, USA) at a rate of 4 ml/min via the renal artery for a total of six hours, followed by a phosphate-buffered saline (PBS) perfusion for 24 hours and 10.0 Ky gamma irradiation. The sterilized scaffolds were then kept at 4 °C.

### Preparation of *in vivo* nuclear staining dyes

The nucleic acid stain Hoechst 33342 (Invitrogen, Carlsbad, CA, USA) was used to image cellular nuclei. Similarly, this infusate was prepared by diluting approximately 50 μl of this cell-permeant probe in 0.5 ml sterilized saline.

### Preparation of large molecular weight fluorescent dextrans infusates

150-kDa fluorescein isothiocyanate (FITC)-dextrans (TdB Consultancy, Uppsala, Sweden), were used to visualize the microvasculature of the native and decellularized kidneys. A stock solution of this dye was first prepared by mixing 20 mg in 1 ml of sterilized saline. Then the infusate was prepared by diluting 500 μl of the stock solution in 1 ml of sterilized saline.

### Intravital imaging of native and transplanted decellularized kidneys using a multiphoton microscope

After anesthetizing each animal, its thoracic region was again shaved, and the topical antiseptic was applied to this area. The animal’s tail vein was then dilated by either placing it in a warm bath or soaking it with a warmed and damped gauze sheet. After identifying the dilated vein, a 25-gauge butterfly needle (Sigma-Aldrich, St. Louis, MO, USA) was inserted into this vascular track to facilitate the infusion of a bolus of 0.5 mL heparinized saline.

Each rat was then transplanted with a scaffold created from its nephrectomized kidney. For this process, flank incisions were then made, and orthotropic transplantations of the biocompatible acellular scaffolds were achieved by performing end-to-end anastomoses of decellularized renal artery and vein to the rat’s previously ligated renal artery and vein, respectively. Anastomoses were performed by occluding the ligated ends of the vessels with micro-serrefine vascular clamps (Fine Science Tools, Foster City, CA, USA) and connected the respective opened ends of the scaffold decellularized vessels with 10-0 silk sutures (Fine Science Tools, Foster City, CA, USA). Fluorescent markers were then infused through the tail vein, and the venous vascular clamp was first removed and then the arterial vascular clamp to reperfuse the scaffold. The sites of anastomoses were carefully inspected and additional sutures were added to eliminate any bleeding that was observed.

The native and transplanted organs were exteriorized for imaging at different measurement time points, as shown in Fig. 1b. Exteriorized native or acellular kidneys were individually positioned inside a 50 mm glass-bottom dish (Willco Wells B.V., Amsterdam, The Netherlands) containing saline and a heating pad was put over the animal to maintain body temperature^28^. Images were then obtained with an Olympus FV1000MP multiphoton/intravital microscope (Center Valley, PA, USA) with a Spectra-Physics MaiTai Deep See laser (Santa Clara, CA, USA), Fig. 1b. The laser was tuned to wavelengths ranging from 770 to 860 nm, which are known to excite the fluorescent dyes used in this study. Images were captured using an X60 water-immersion objective and external blue- and green-based emission detectors.

### Biochemical assay to measure decellularization efficacies

Using a FastPrep 24 Tissue Homogenizer (MP Biochemicals, Santa Ana, CA, USA) the acellular whole organs were homogenized, and 1 ml methylene blue solution (methylene blue 0.25 g/l, anhydrous sodium sulfate 50 g/l, concentrated sulfuric acid 10 ml/l) was added to the suspension. The suspension was then digested at 56 °C with proteinase K (Omega Bio-tek, Atlanta, GA, USA) for approximately 1 hour. The samples were then isolated with chloroform, and the SDS content of the extracts was determined using a Molecular Devices SpectraMax M Series Multi-Mode Microplate Reader (Sunnyvale, CA, USA) to obtain absorbance measurements at a wavelength of 650 nm. Concurrently, DNA contents were also quantified Using a Qiagen DNeasy Kit (Qiagen, Valencia, CA, USA) by first storing samples of the minced sections at −80°C overnight. Thereafter, the tissues were lyophilized and the ratios of ng DNA per mg dry tissue were estimated using an Invitrogen Quant-iT PicoGreen dsDNA assay kit (Carlsbad, CA, USA) and the microplate reader.

### Intravital microscopy assay to measure decellularization efficacies

Static micrographs and time-series videos were obtained using the multi-photon microscope system. Imaging data was collected on day 0, day 1, day 2, and day 7 time points to examine cell nuclear patterns in native kidneys and scaffolds, changes in innate/decellularized tissue fluorescence levels, and microvascular integrity/function, using well-established techniques^13,15,18–24,26–29–31^. To conduct this analysis, we randomly selected 4 regions within a microscopic field and measured the blue and green pseudocolor fluorescence levels in nuclear and epithelial compartments, respectively.

### Assessments of structural/functional integrities and microcirculation velocities in native and decellularized nephrons

Additional qualitative and quantitative assessments of microcirculation velocity were obtained by tracking the linear displacement of blood cells within a given period. Finally, microarchitecture porosity and tubular/glomerular integrity estimations were acquired by calculating extravasation rates from various nephron compartments. These rates were obtained by measuring the quantities of the fluorescent dextran that leaked from various capillary vasculature and entered the tubular epithelium and lumen.

### Data Records

The data is located on a publicly available data repository, called Organ Scaffold Characterizations Using Intravital Microscopy Repository, with the following Digital Object Identifier (DOI): https://doi.org/10.6084/m9.figshare.18703949.

### Technical Validation

Several measures were taken to ensure reliability and that the collected data was unbiased. Rats were assigned at random and housed in the same cage. They were all the same breed and were provided with the same feeding conditions. Additionally, rats were checked for signs of illness or infection before all experiments. All animal surgical procedures and decellularization techniques were conducted by the same researcher. Also, all and IVM studies were conducted by the same researcher using the same microscope settings. Upon curation of the data, all biochemical data were logged in an Excel file for analysis. Olympus OIF data were analyzed using FV10-ASW Viewer 3.1 (Olympus corporation, Shinjuku, Tokyo, Japan) and ImageJ (Fiji-ImageJ 64, US National Institutes of Health, Bethesda, MD, USA) software packages. Time series data files are listed in the data collection to easily highlight intravital events capturing events within the transplanted scaffolds. We reviewed all the logged data to ensure that no data entry errors occurred before our analyses. As a negative control, decellularized samples were infused with saline solution. No nuclear staining or vascular lining was observed, as expected.

### Biochemical estimations of DNA and SDS contents in native and decellularized kidneys

The results from these procedures are presented in Fig. 2. These results revealed that the substantial visible changes to the native organ correlated with the removal of approximately 98% of the innate DNA content from original kidneys (Kruskal-Wallis, H = 23.298, d.f. = 3, p < 0.001), as previously reported ^11,32^. Moreover, the assays revealed that roughly 99% of the SDS used to facilitate decellularization was removed at the end of the process (Kruskal-Wallis, H = 23.345, d.f. = 3, p < 0.001). These results highlight the efficacy of the decellularization protocol. It is vital to efficiently eliminate cellular remains from scaffolds to limit the risk of undesired immunological responses or tissue rejection following transplantation^23^. A critical feature of the decellularization process is determining the concentrations of these remaining components. It is also critical that residual SDS concentrations are kept low enough to prevent the detergent from further denaturing the scaffold. This undesired action can change the scaffold’s overall permeability and impair its integrity, making recellularization and transplantation more difficult^24^.

**Fig. 2.**
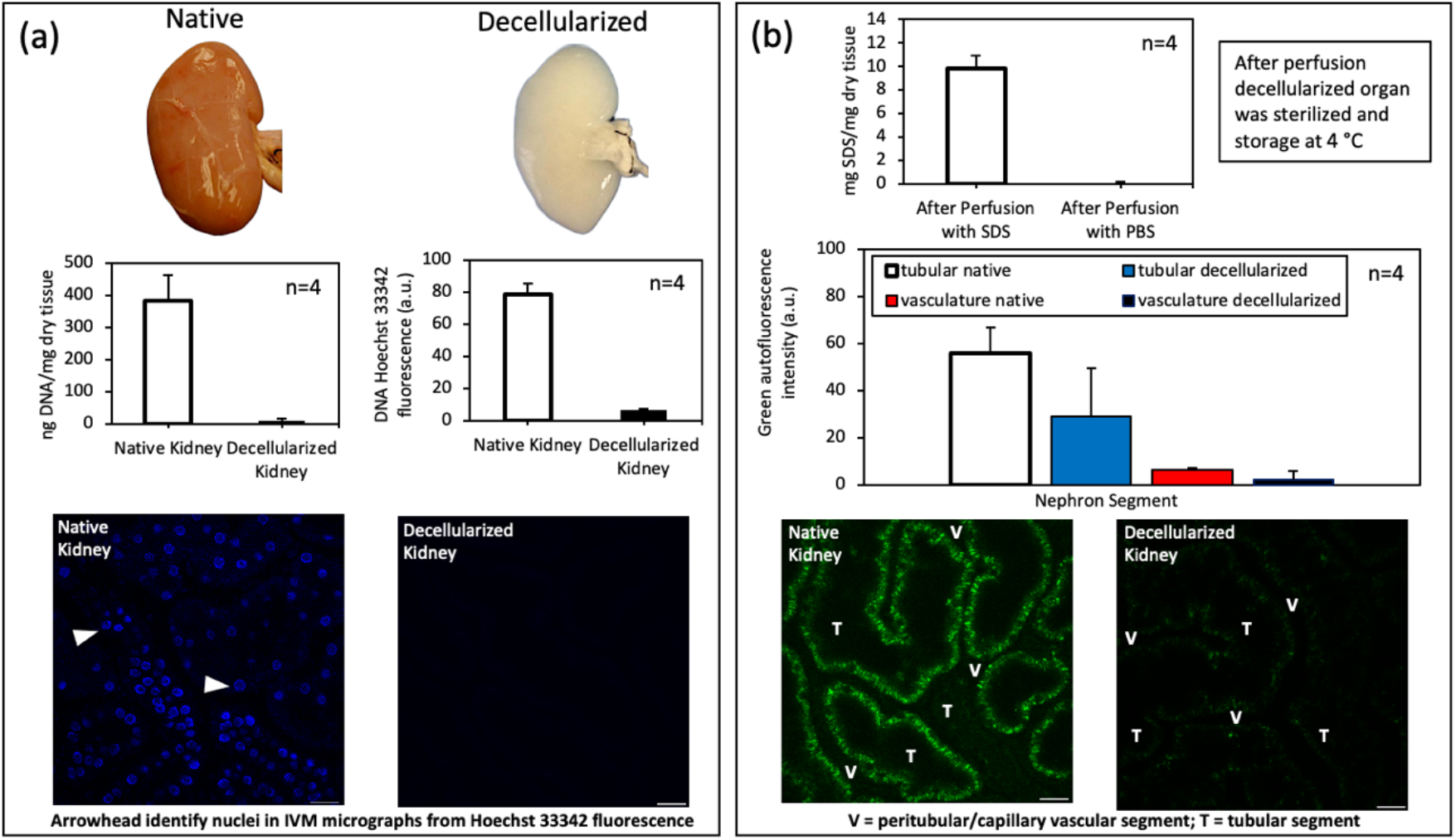
Biochemical and IVM assays that evaluated the decellularization efficacy, (**a**) Graphical displays of DNA concentrations (ng DNA / mg dry tissue) in native and acellular kidneys from biochemical assay correlated with Hoechst 33342 signals (a.u.) in IVM micrographs from fluorescently labeled DNA (arrowheads) in these tissue types, (**b**) Graphical illustrations of SDS removal from scaffolds with PBS and changes in tissue fluorescence with decellularization using IVM micrographs. Kruskal-Wallis tests detected significant DNA, SDS, and fluorescence after decellularization (p < 0.001). Scale bar 20 μm.

**Fig. 3.**
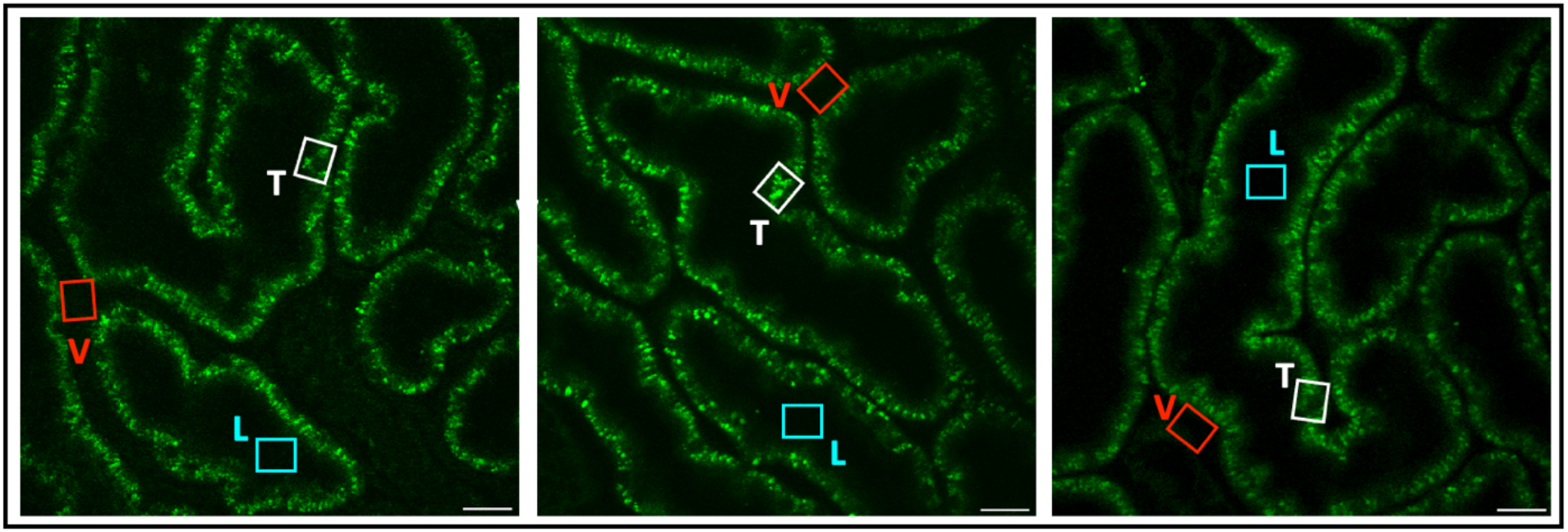
IVM micrographs of native kidneys. These images were analyzed to determine the fluorescence levels within the vasculature, tubular epithelium and tubular lumen labeled V, T, and L, respectively. These micrographs show normal tissue fluorescence patterns within these various nephron segments. The highlighted segments are representative of the regions selected to record the fluorescence intensity using ImageJ software. Scale bar 20 μm.

**Fig. 4.**
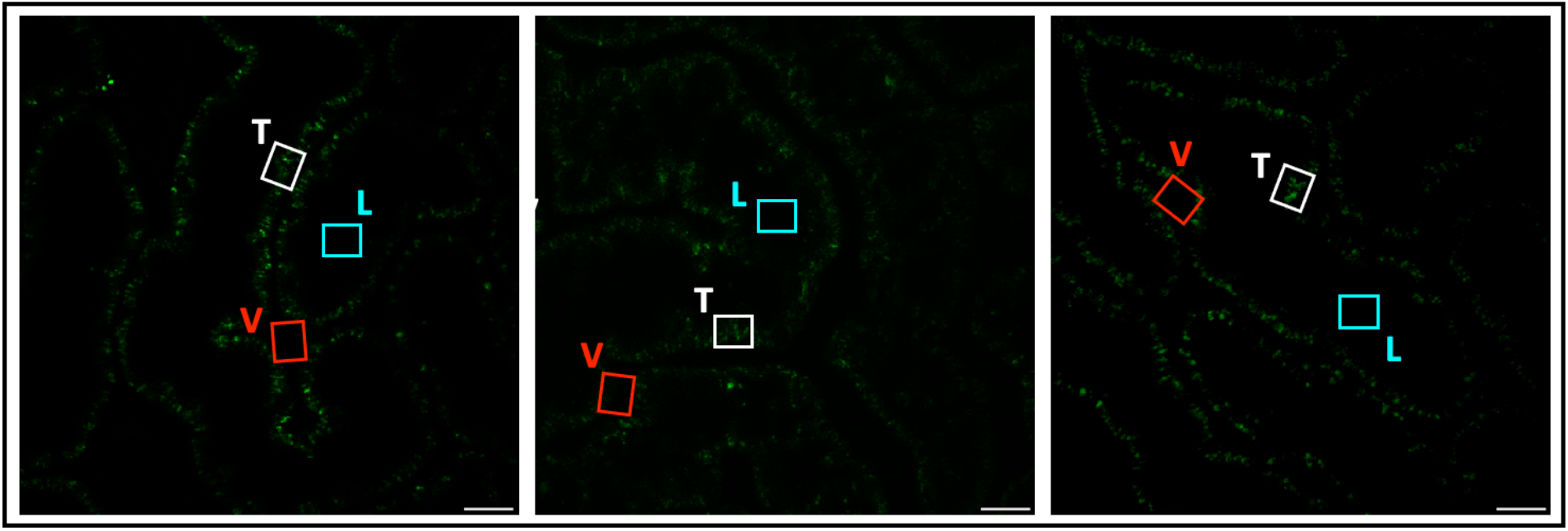
IVM micrographs of decellularized scaffolds Day 0. These images were analyzed to determine the fluorescence levels within the vasculature, tubular epithelium and tubular lumen labeled V, T, and L, respectively. These micrographs show substantial reductions in tissue fluorescence levels within these various nephron segments generated from decellularization to obtain intrinsic fluorescence patterns in the acellular organ. The highlighted segments are representative of the regions selected to record the fluorescence intensity using ImageJ software. Scale bar 20 μm.

**Fig. 5.**
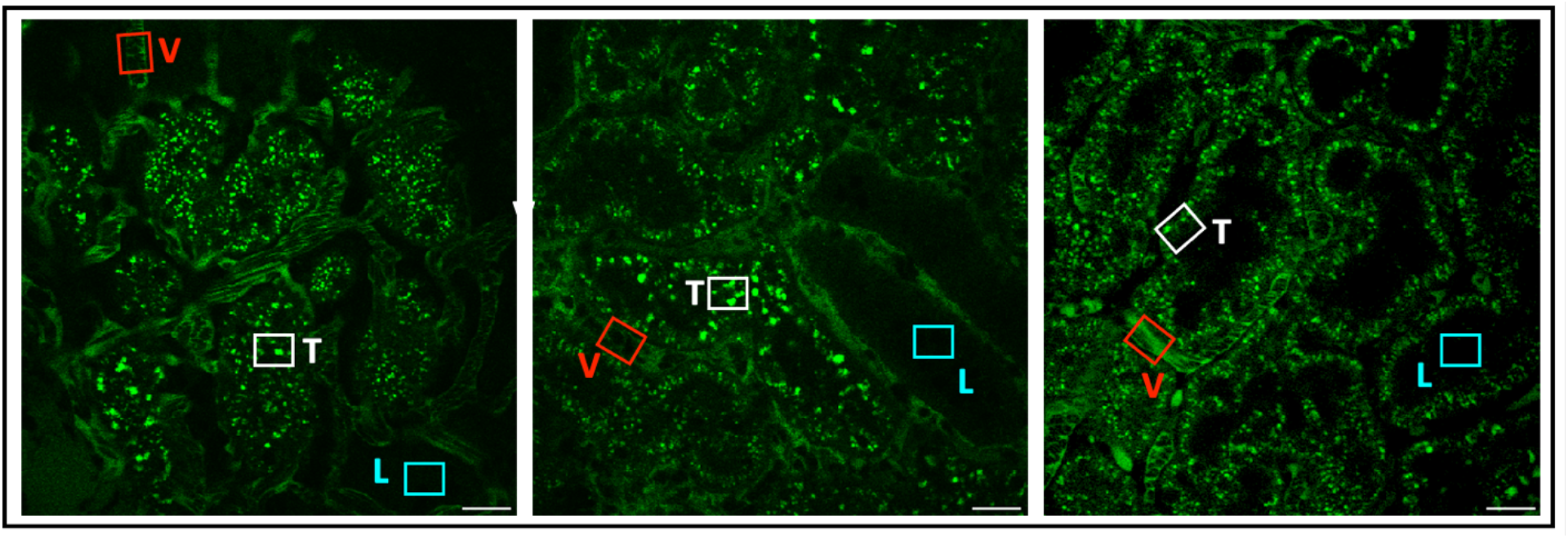
IVM micrographs of decellularized scaffolds Day 1 after transplantation. These images were analyzed to determine the fluorescence levels within the vasculature, tubular epithelium and tubular lumen labeled V, T, and L, respectively. These micrographs show the presence of ischemic blood flow, increases in tubular and luminal fluorescence levels, and damage to the tubular architecture within these various nephron segments generated from decellularization. The highlighted segments are representative of the regions selected to record the fluorescence intensity using ImageJ software. Scale bar 20 μm.

**Fig. 6.**
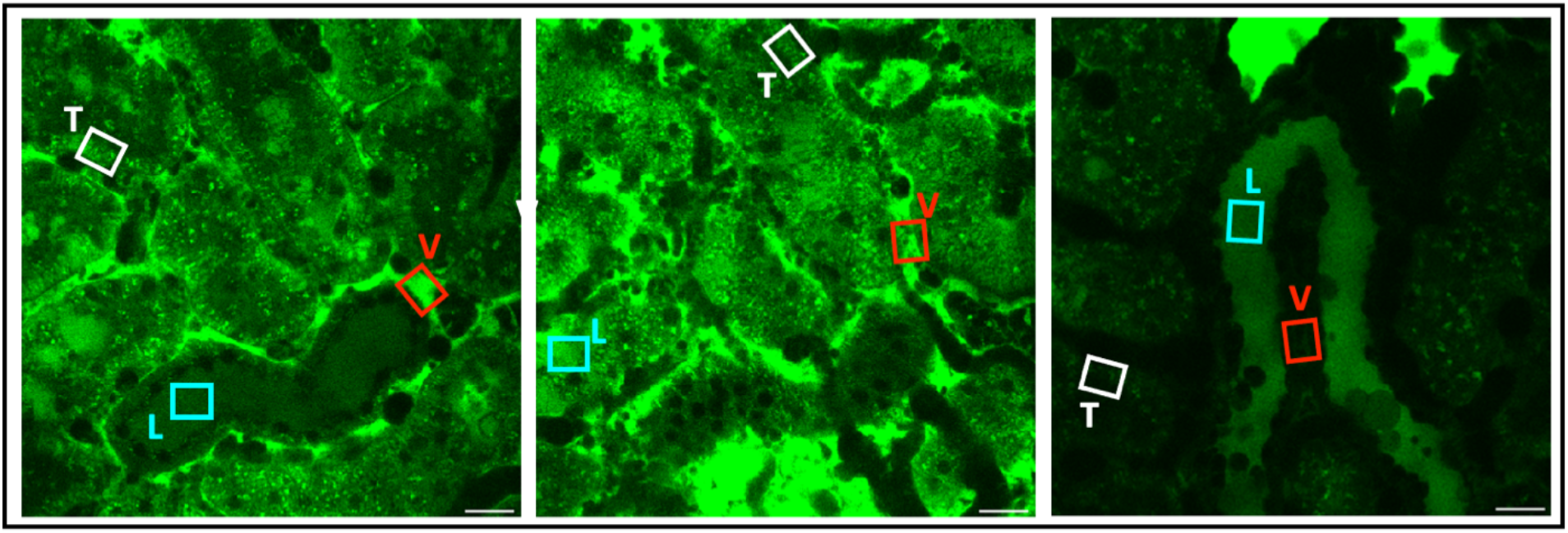
IVM micrographs of decellularized scaffolds Day 2 after transplantation. These images were analyzed to determine the fluorescence levels within the vasculature, tubular epithelium and tubular lumen labeled V, T, and L, respectively. These micrographs show substantial levels of ischemia, dextran leakage from the decellularized vasculature, and bleb formation with decellularized tubular lumens. The highlighted segments are representative of the regions selected to record the fluorescence intensity using ImageJ software. Scale bar 20 μm.

**Fig. 7.**
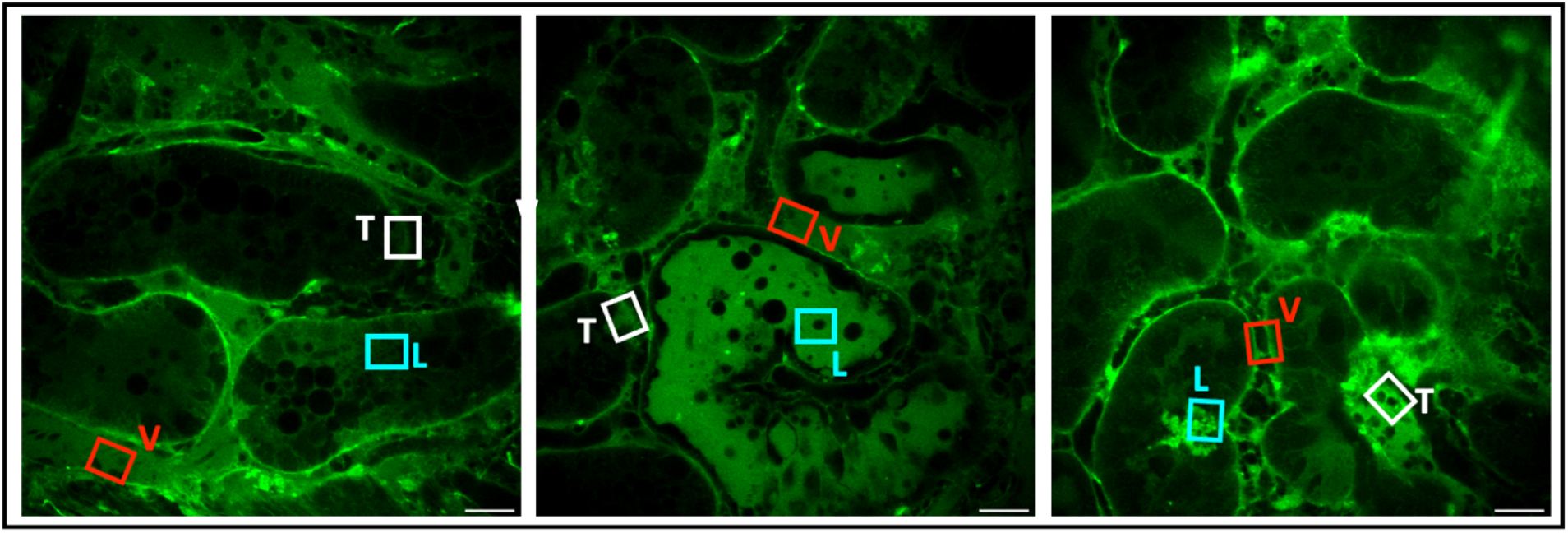
IVM micrographs of decellularized scaffolds Day 7 after transplantation. These images were analyzed to determine the fluorescence levels within the vasculature, tubular epithelium and tubular lumen labeled V, T, and L, respectively. The highlighted segments are representative of the regions selected to record the fluorescence intensity using ImageJ software. These micrographs again show substantial levels of ischemia, dextran leakage from the decellularized vasculature, and bleb formation with decellularized tubular lumens that supported the progressive demise of the transplanted scaffolds. Scale bar 20 μm.

**Fig. 8.**
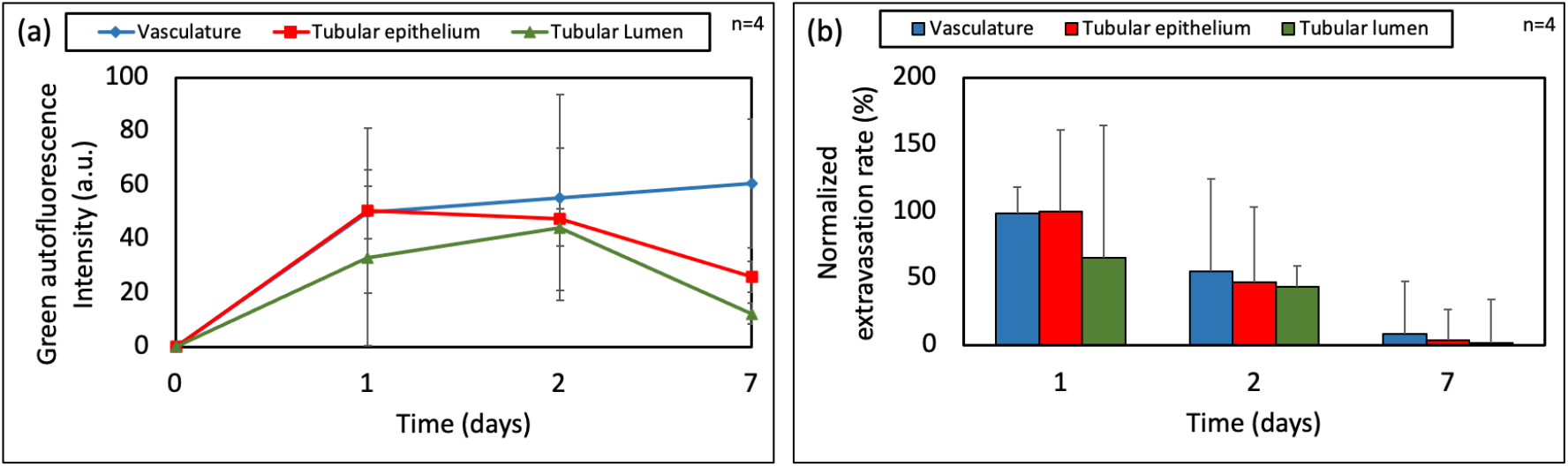
Loss of structural and functional integrities of various decellularized nephron segments after transplantation, (**a**) Estimations of microarchitecture porosity obtained using IVM reveal substantial increases in fluorescence intensity signals, highlighting the progressive presence of 150-kDa fluorescein isothiocyanate (FITC)-dextran molecules within various decellularized nephron components, (**b**) IVM micrographs were used to track degrees of extensive blood extravasation from the decellularized vasculature into peritubular and tubular compartments that occurred during the 7 days post-transplantation.

### IVM estimations of DNA contents in native and decellularized kidneys

Qualitative and quantitative assessments of tissue fluorescence patterns were conducted to evaluate the effect of the decellularization process and determine its ability to characterize the conversion of native kidneys into acellular structures adequately (Fig 2). Several measures were taken to ensure reliability and that the collected data was unbiased. Rats were collected at random and housed in the same cage. They were all the same breed and were provided with the same feeding conditions. Additionally, rats were checked for signs of illness or infection before experimentation, and after surgical procedures and imaging sessions were conducted. Common negative controls used in IVM studies were applied to confirm the nature of the tissue fluorescence in native and decellularized kidneys.

The samples were infused with saline solution and no dye as a negative control during the imaging process. As expected, no nuclear staining was observed in the negative control; we were only able to observe the auto fluorescent background. IVM was used to further investigate the efficiency of the decellularization process as a proof of principle. The fluorescence of native/non-transplanted kidneys and decellularized scaffolds that were transplanted into living patients after introducing Hoechst 33342 exhibited significant differences in fluorescence. This membrane-permeant dye easily penetrates tissues^25–30^, where it identifies DNA in living and fixed cells by binding to adenine-thymine-rich sections of DNA in the minor groove, resulting in substantial fluorescence enhancements^31^. The clear imaging of nuclei in endothelia, epithelia, glomeruli, interstitial cells, and circulating leukocytes is known to benefit from such enhancements^28^.

In comparison to the native kidney, fluorescence intensity measurements revealed a 92 percent reduction in the relative amount of fluorescence in the decellularized scaffold and the lack of nuclear staining. This drop in relative blue pseudo-color fluorescence corresponds to a decrease in DNA content as determined by the biochemical experiment. These *in vivo* findings are in addition to the standard *in vitro* histological and fluorescence microscopic approaches used to evaluate decellularization^32^.

Imaging conducted with the green pseudo-color channel revealed differences in the intrinsic frequency of auto fluorescence in native kidneys compared to the decellularized organ. The kidney normally has a high level of green autofluorescence^25, 26^, which is caused by biological structures such as mitochondria and lysosomes^20^, as well as unique metabolites such as aromatic flavins, nicotinamide adenine dinucleotide and amino acids^33, 34^, emitting light naturally. Without the use of fluorescent markers, the relative distribution and percentages of these structures assist differentiate renal compartments, such as the proximal and distal tubules, glomerulus, peritubular capillaries, and interstitium^9, 28^.

Proximal tubules have the strongest autofluorescence signal and thickness, whereas distal tubules seem thinner and dimmer. Similarly, the Bowman’s capsule contour and faint or nonexistent capillary tuft^35^, as well as the surrounding peritubular capillary and interstitial space, indicate the renal corpuscle’s distinctive shape. Decellularization, on the other hand, resulted in significant reductions in the green pseudo-color signal intensity. The relative level of fluorescence in the scaffolds dropped by about half, making it impossible to distinguish various tubular compartments, with the exception of proximal tubule segments emanating from the decellularized Bowman’s capsule. Using immunohistochemical techniques *in vitro*, similar significant reductions in tissue fluorescence were previously documented, confirming decellularization^12, 24, 36^.

### Blood extravasation rates and changes in microarchitecture porosity and tubular/glomerular integrity

During the week following transplantation, vascular permeability was further studied. Average extravasation rates were calculated across the 7-day experimental period. The average extravasation rate was calculated by taking the mean fluorescence value for each time point for each renal compartment and dividing that by the number of hours that had elapsed after transplantation. Reductions in the average extravasation rates within various decellularized nephron segments were observed after the 150-kDa FITC dextran was infused into the tail vein before blood was introduced into the transplanted scaffolds. This method allowed us to assess the distribution of blood contents as well as get insight into the effect of the *in vivo* environment on the grafts. This infusion regimen was chosen over later infusions because it was vital to ensure that the fluorescent marker circulated through the nephron since clotting could prevent the dye from reaching the microvasculature at later time points.

Compared to the initial state, substantial alterations in vascular structure and function were found at the Day 1, Day 2, and Day 7 timepoints. Under *in vivo* settings, the variable degrees of dextran extravasation from the microvasculature indicate impairment of normal filtrative capacities and increased permeability of the decellularized microvasculature. The kidney may normally autoregulate blood flow to protect peritubular and glomerular capillaries from significant variations in blood pressure^37^. The gradual leakage of the FITC dye from the glomerular capillaries into the Bowman’s space may be explained by such damage. Decellularization would have advertently impacted microarchitectural permeability, impairing the glomerulus’ ability to operate as a natural sieve, allowing only water and tiny solutes to pass into the filtrate. Decellularization removed this intrinsic semipermeable barrier, enabling macromolecular transport to be unrestricted by size, shape, charge, or deformability^39^.

### *In vivo* assessment of velocity within the microcirculation of native kidneys and transplanted acellular scaffolds

We estimated the velocity within the microcirculation of native kidneys and transplanted acellular scaffolds using times-series IVM data. The velocities were computed from the ratios of blood displacement and time. The video sequences were collected to support these estimations and the normalized blood flow rates within the peritubular capillaries of native (non-transplanted) kidney were compared with the velocities in transplanted scaffolds after transplantation at Day 0, Day 1, Day 2, and Day7. Static images are presented in Fig. 9a outlining the microscopic movement within various vascular tracks and timestamps that were used to measure duration. The Kruskal-Wallis test detected a significant difference among velocities within microcirculatory systems in the abovementioned cases (p < 0.019), and Dunn’s post hoc test revealed pairwise differences between the velocity of the microcirculation of native kidneys and scaffolds on day 2 (p = 0.002), scaffolds on day 0 and scaffolds on day 2 (p = 0.012), scaffolds on day 20 and scaffolds on day 7 (p = 0.019). These results can be found in Fig. 9.

**Fig. 9.**
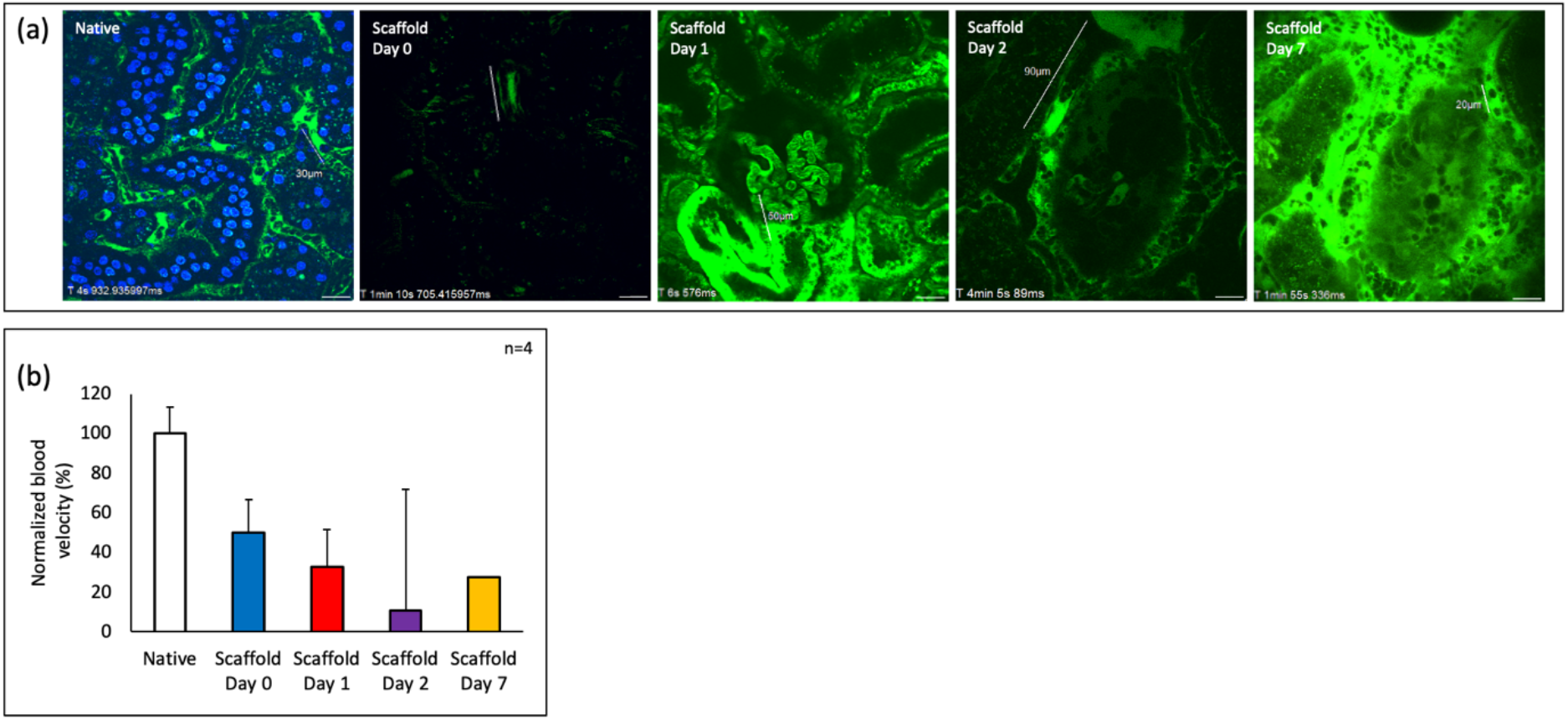
In vivo assessment of velocity within the microcirculation of native kidneys and transplanted acellular scaffolds, (**a**) IVM micrographs were taken to estimate blood flow rates in the peritubular capillaries of native (non-transplanted) kidneys and transplanted scaffolds directly after transplantation at Day 0, Day 1, Day 2, and Day7. (**b**) Kruskal-Wallis tests detected a significant difference among velocities within microcirculatory systems in the abovementioned cases (p < 0.019), and pairwise differences were observed between the velocity of the microcirculation of native kidneys and scaffolds on day 2 (p = 0.002), scaffolds on day 0 and scaffolds on day 2 (p = 0.012), scaffolds on day 20 and scaffolds on day 7 (p = 0.019).

### Usage Notes

Data processing and analysis procedures, and the results for biochemical assays (UV-visible spectroscopy DNA and SDS concentration measurements), and IVM assays (DNA, cell, and tissue fluorescent intensity measurements, microvascular leakage, tubular and glomerular integrity, and renal capillary blood velocity) are presented in our repository in the file called Organ Scaffold Characterizations Using Intravital Microscopy Images.xlsx. This file outlines specific details on the ways we handled the data. Time series IVM and static images files are presented in the folder used to measure velocity in the microcirculation. Our group and others in the field have previously used biochemical assays to analyze decellularization efficacy. Moreover, IVM analyses performed have been adapted from related conducted by our group and several others. In both cases, we have outlined the respective citations. The entire data is publicly available under a Creative Commons CCO license to support its use for further analysis and publication under the requirement of citing this article and the dataset.

## Acknowledgments

The project was supported by an Institutional Research and Academic Career Development Award (IRACDA), Grant Number: NIH/NIGMS K12-GM102773, and support from Khalifa University of Science and Technology, Grant Numbers: FSU-2020-25 and RC2-2018-022 (HEIC). We thank Dr. Joao Paulo Zambon, Dr. Amanda Dillard, and Mr. Kenneth Grant for their support in developing the experimental models. We also thank Mrs. Maja Corridon and Ms. Xinyu Wang for reviewing the manuscript.

## Contributions

Peter R. Corridon: Conception and design of the study, data collection supervision, design of data sharing plan, revision, data preparation, validation, quality control, uploading, manuscript figure making, manuscript writing, revision, and final approval of the version submitted for publication. Anousha A. Khan: data preparation, validation, manuscript figure making, manuscript writing, review, and final approval of the version submitted for publication.

